# Characterization of directed differentiation by high-throughput single-cell RNA-Seq

**DOI:** 10.1101/003236

**Authors:** Magali Soumillon, Davide Cacchiarelli, Stefan Semrau, Alexander van Oudenaarden, Tarjei S. Mikkelsen

## Abstract

Directed differentiation of cells *in vitro* is a powerful approach for dissection of developmental pathways, disease modeling and regenerative medicine, but analysis of such systems is complicated by heterogeneous and asynchronous cellular responses to differentiation-inducing stimuli. To enable deep characterization of heterogeneous cell populations, we developed an efficient digital gene expression profiling protocol that enables surveying of mRNA in thousands of single cells at a time. We then applied this protocol to profile 12,832 cells collected at multiple time points during directed adipogenic differentiation of human adipose-derived stem/stromal cells *in vitro*. The resulting data reveal the major axes of cell-to-cell variation within and between time points, and an inverse relationship between inflammatory gene expression and lipid accumulation across cells from a single donor.

---

Single-cell transcriptome profiling enabled by next-generation sequencing has recently emerged as a promising tool for characterization of heterogeneous cell populations^1^, but routine adoption will require development of protocols that allow unbiased profiling of large numbers of cells at reasonable cost. To enable efficient characterization of thousands of single cells at a time, we developed a 3′ digital gene expression (3′ DGE) RNA-Seq protocol that we refer to as SCRB-Seq (single cell RNA barcoding and sequencing; Supplementary Methods, Figure S1 and Table S1). Building on recent advances^2–6^, SCRB-Seq relies on a template-switching reverse transcriptase to convert poly(A)+ mRNA from isolated single cells to cDNA decorated with universal adapters, well-specific barcodes and unique molecular identifiers (UMIs)^7^. Decorated cDNA from multiple cells are then pooled, amplified and prepared for multiplexed sequencing using a modified transposon-based fragmentation approach that enriches for 3′ ends and preserves strand information. SCRB-Seq is specifically optimized for surveying mRNA from large cell numbers using minimal reagents, reagent transfers and sequencing reads per cell, with the aim of characterizing the major patterns of gene expression variation across heterogeneous populations in a cost-efficient manner. It requires approximately two times fewer enzymatic reactions, purifications and liquid transfer steps than a previous high-throughput protocol^8^ and is complementary to protocols that are optimized for deep, full-length transcriptome coverage^9^.

To demonstrate SCRB-Seq, we applied it to characterize a primary human adipose-derived stem/stromal cell (hASC)^10^ differentiation system (see Supplementary Materials and Figure S2). *In vitro* adipogenesis is both a general model of lineage commitment and an important source of cells for research on metabolic disorders^11,12^. A variety of cell populations can be induced to differentiate by cocktails of adipogenic hormones and growth factors, but their yields of lipid-filled, adipocyte-like cells are highly variable. It remains unclear to what extent this variability reflects heterogeneity in the starting populations, stochastic responses to imperfect differentiation stimuli or other factors. The great majority of adipogenesis research over the last three decades has therefore relied on the immortalized murine 3T3-L1 cell line, which supports near complete conversion to adipocyte-like cells^13^. Numerous molecular differences have, however, been found between this cell line and hASCs^14^. Single-cell profiling should help clarify the origin and relevance of these differences and also improve the utility of more heterogeneous differentiation systems.

We collected cells from hASC cultures just prior to induction of differentiation (day 0), as well as at seven time points after induction (days 1, 2, 3, 5, 7, 9 and 14). At the last time point, approximately two thirds of the cells contained clearly visible lipid droplets while the remainder retained a more fibroblast-like morphology (Figure 1A). We used a nucleic acid stain to identify and sort intact single cells into 384-well plates with a fluorescence-activated cell sorter (FACS; Supplementary Figure 2A). We also used a neutral lipid stain to separately sort single cells based on their lipid contents (Supplementary Figure S2B). After further protocol optimizations (see Supplementary Methods), we collected and sorted additional cells from independent cultures at days 0, 3 and 7 (Supplementary Figures S2 and S3). In total, we prepared multiplexed SCBR-Seq libraries from 44 microplates (Supplementary Tables S1 and S2), sequenced these to a mean depth of ∼165,000 reads per well and then aligned the reads to RefSeq transcripts. After stringent filtering on sequence and alignment quality, and then estimating the expression levels in each cell from UMI counts (Supplementary Figure S4), we obtained survey-depth DGE profiles from a total of 12,832 cells (76% of the total wells). As judged by the UMI counts, each DGE profile captured between 1,000 and ∼10,000 unique mRNAs (mean = 2,602 and 3,336 for the initial and optimized protocols, respectively), which constitutes a ∼4-fold increase in mean library complexity relative to a previous high-throughput protocol^8^.

**Figure 1.**
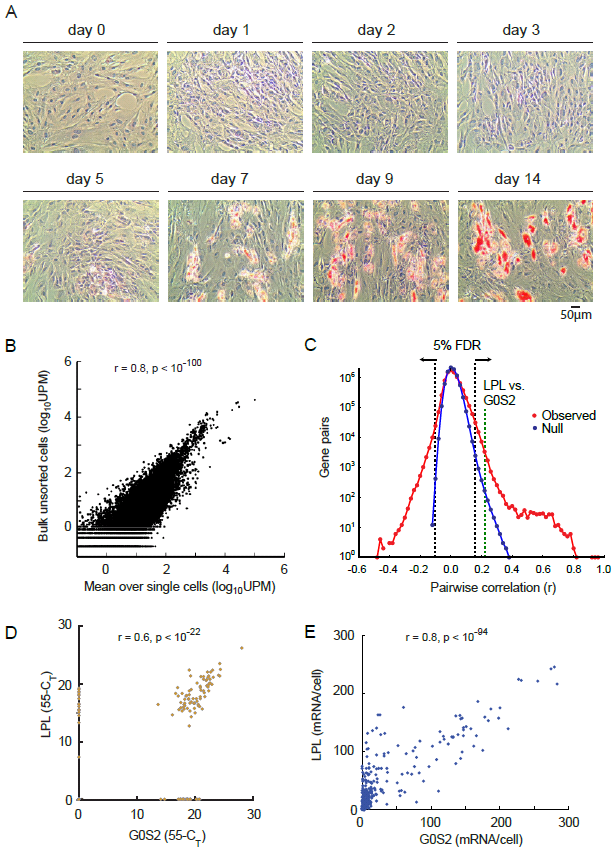
Characterization of cell culture heterogeneity using SCRB-Seq. **A)** Heterogeneity in hASC cultures upon induction of directed adipogenesis. Lipid accumulation is visualized by Oil Red O staining. **B)** Gene expression estimates from bulk cells compared to their corresponding means across the single cell profiles. UPM = UMI counts for one gene per million UMI counts for all genes. **C)** Distribution of observed pairwise correlations (Pearson’s r) between all pairs of genes that were detected in at least 10% of day 7 cells (n = 4,038 genes) compared to an estimated null distribution obtained by permuting the expression values of each gene across the same cells. **D)** Single cell qRT-PCR and **E)** smFISH validation of the observed positive correlation between LPL and G0S2 from separate cells also collected at day 7.

**Figure 2.**
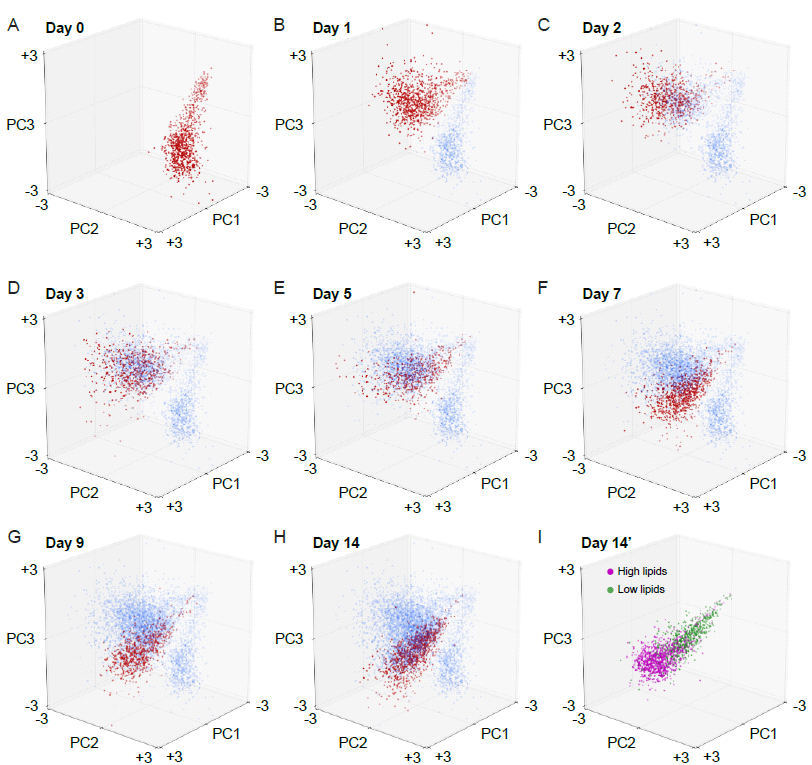
Gene expression dynamics at single cell resolution. **A-H)** Each scatter plot shows the first three PCs of the initial hASC time course. Red dots show cells collected at the indicated time point, while blue dots show cells collected at all previous time points. **I)** Separately sorted cells with high and low lipid content from day 14 projected into the same PC space.

Initial analysis of the resulting data showed that the mean gene expression levels across the single cell profiles were significantly correlated with their corresponding levels from bulk unsorted cells collected at the same time point (r = 0.8, p < 10^−100^; Figure 1B). Of 15,099 distinct RefSeq genes that were detected at day 0 in bulk unsorted cells, 14,612 (97%) were also detected in at least one single cell from the same day. As expected from the relatively low sequencing coverage, only the most actively transcribed genes were captured from every cell (Supplementary Figure S5), but we could nevertheless detect significant positive and negative correlations between the expression levels of individual genes across cells collected on the same day (Figure 1C; Supplementary Figure S6). For example, LPL and G0S2, two traditional markers that are both up-regulated after induction of adipogenesis, had positively correlated expression levels after differentiation (r = 0.23, p < 10^−12^ on day 7; FDR ≤ 5%). We could validate a positive correlation between these genes both by qRT-PCR analysis of independently sorted single cells (Figure 1D) and *in situ* by multiplexed single molecule FISH^15^ (smFISH; Figure 1E; Supplementary Figure S7 and Table S3). Further comparison suggested that the mRNA detection efficiency of SCRB-Seq at the chosen sequencing depth was approximately 1-2% relative to smFISH (LPL: mean un-normalized UMI count = 1.0/cell, range = 0 to 18 from SCRB-Seq, mean mRNA count = 44.7/cell, range = 0 to 350 from smFISH; G0S2: mean un-normalized UMI count = 0.26/cell, range = 0 to 10 from SCRB-Seq, mean mRNA count = 31.2, range = 0 to 313 from smFISH). We conclude that SCRB-Seq can capture gene expression variation at single-cell resolution, although the relatively unbiased transcriptome coverage comes at the cost of lower sensitivity than state-of-the-art single molecule detection methods.

To understand the observed cell-to-cell variation in gene expression in more detail, we performed a principal component analysis (PCA) of the initial time course (days 0 to 14; 6,197 cells; Figure 2A-H). Plotting the position of each cell in the space defined by the first three principal components revealed two salient features. First, there was little overlap between cells from day 0 and cells from later time points, which suggests that addition of the adipogenic differentiation cocktail induced a rapid response in virtually all of the cultured cells. Second, gene expression levels continued to evolve from day 1 to day 14, but there was substantial overlap between the cells collected at close time points. This is consistent with a population-wide, but asynchronous, response to induction of differentiation.

To explore the biological basis for the observed gene expression variation, we next examined the relationships between each of the top principal components (PCs), gene expression and time (Figure 3). The PCs can be interpreted as metagenes^16^ that capture coordinated expression of multiple genes in the original data set. For each PC, we therefore ranked the genes according to their corresponding PC weights and then looked for evidence of coordinately regulated pathways using gene set enrichment analysis^17^ (GSEA; Supplementary Data S1). This analysis suggested qualitative biological interpretations for at least the top four PCs.

**Figure 3.**
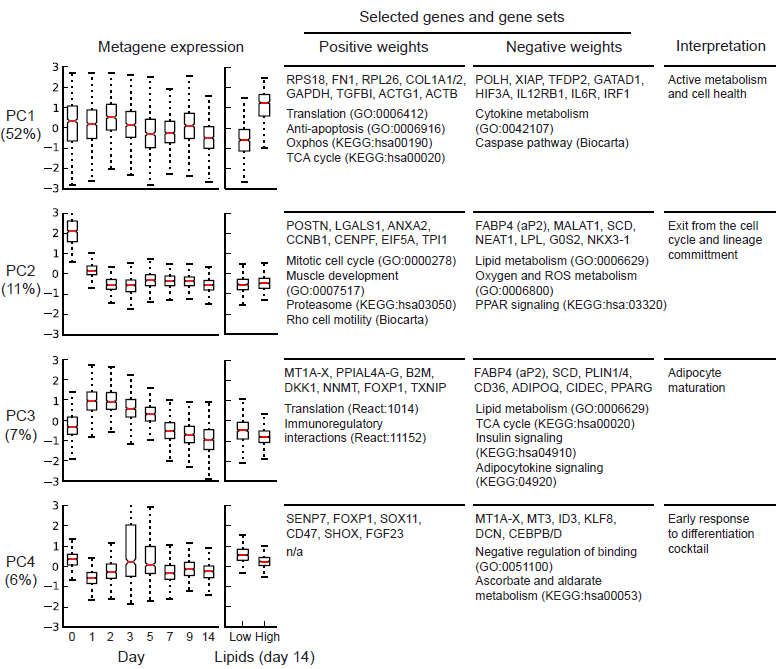
Gene set enrichment analysis. Distributions of weights (right) and selected genes and genes sets associated with positive and negative weights (left) for the top four PCs in the initial hASC time course and lipid-based sort. See also Supplementary Data S1-S4 for complete GSEA results. Percentages indicate the ratio of the total variance in the data set captured by each PC. Red lines indicate medians, boxes the 1^st^ and 3^rd^ quartiles and whiskers the ranges.

The first PC metagene (PC1) was positively associated with genes involved in general cellular metabolism, including the majority of genes involved in ribosome assembly, mitochondrial biogenesis and oxidative phosphorylation, while it was negatively associated with inflammatory pathways, cytokine production and caspase expression. We interpret variation along PC1 to reflect differences between metabolically active “healthy” and inactive “unhealthy” cells. Interestingly, while there was a shift towards the latter state towards day 14, there was substantial overlap between the PC1 distributions from all time points, which indicates that this axis of variation was a major contributor to culture heterogeneity prior to induction of differentiation. Note that we did not observe significant cell detachment or death during the two weeks of differentiation, which suggests that the inflammation signature represents a chronic cell state rather than ongoing apoptosis. In contrast, PC2 was high only in cells collected from day 0, effectively separating these from the differentiating cells. It showed a strong positive association with expression of genes required for progression through the mitotic cell cycle and to a lesser extent with genes associated with non-adipogenic differentiation. We therefore interpret a decrease in PC2 to reflect exit from the cell cycle and lineage commitment. Expression of PC3 was high during the first two days post-induction and then steadily decreased towards day 14. This decrease was associated with up-regulation of lipid homeostasis pathways and markers of adipocyte maturation. PC4 showed a transient drop at day 1, which was associated with increased expression of genes known to be rapidly induced by adipogenic cocktails, including early adipogenic regulators CEBPB and CEBPD^11^. We therefore interpret PC4 to reflect an early response to induction of differentiation.

To explore the relationship between variations in gene expression and in lipid droplet accumulation, we next analyzed an additional 933 cells with high and 666 cells with low lipid contents collected at day 14 (Supplementary Figure S2). When we projected the DGE profiles of these cells into the space defined by the initial time course PCs, we found that high and low lipid cells were largely separated by their distribution along PC1 (Figure 2I, Figure 3). That is, cells with higher lipid content showed higher expression of genes related to basic cellular metabolism, while cells with lower lipid content showed higher expression of inflammatory genes. Interestingly, there was substantial overlap along PC3, and while some classic adipocyte markers like FABP4 (aP2) were enriched in the high lipid fraction (Supplementary Data S2), key regulatory factors such as PPARG were not, which implies that pathways related to lipid homeostasis and adipocyte maturation had been activated in both fractions. Separate PCAs of the second collected time course (2,968 cells from days 0, 3 and 7, and 2,068 additional cells with high or low lipids from day 7) yielded qualitatively similar patterns (Supplementary Figure S8 and Data S3 and S4), which suggests that our observations are robust to technical variation across cell cultures.

Thus, while morphological analysis suggested that only a fraction of hASCs respond to the differentiation cocktail, our single-cell data show that virtually all of the cells exited the mitotic cell cycle and proceeded to up-regulate an adipogenic gene expression program. The observed variability in lipid droplet accumulation and conversion to mature adipocyte-like morphologies is instead most strongly linked to an inverse correlation in expression of basic cellular metabolism and inflammatory expression programs, which was also present prior to the induction of differentiation. Notably, cells with low lipid contents showed elevated expression of several pro-inflammatory regulatory factors, including as IRF1, IRF3 and IRF4 (Supplementary Data S2 and S4). These factors have previously been shown to negatively influence total lipid accumulation in murine bulk cultures and in vivo models^18,19^, which support a causal link between cell-to-cell variation in expression of these factors and lipid accumulation. Specific activation in the fraction of low lipid cells may explain the paradoxical increases in expression of these factors that have previously been observed in bulk cultures^18^. Interestingly, increased inflammatory expression and decreased metabolism has also been observed in bulk profiling of lipoaspirates from human subjects with adipose tissue dysfunction^20–22^. This increase in part reflects immune infiltration, but also activation of inflammatory cascades in adipocytes^23^. This parallel variation in gene expression between single cells from the same donor and between tissues from multiple donors merits further study.

In conclusion, we have applied SCRB-Seq to survey gene expression in differentiating hASC cultures at single cell resolution. The resulting data reveal the major axes of variation on gene expression, suggest a biological basis for the morphological heterogeneity observed in these cultures, and provide a rich resource for dissection of the regulatory networks involved in adipocyte formation and function. Future advances in sequencing and cell isolation technologies will enable identification of rare expression programs through deeper and more sensitive profiling of every cell, and direct comparison of in vitro and in vivo heterogeneity through direct profiling of single cells from tissue samples.

## Acknowledgements

The authors are grateful to S. Ionescu, J. LaVecchio, C. Trapnell, X. Adiconis, J. Levin, A. Shalek, N. Ruderman and E. D. Rosen for helpful discussions, the staff of HSCRB-HSCI Flow Cytometry Core for assistance with FACS and the staff of the Bauer Core facilities for assistance with sequencing. This work was supported by funds from the Harvard Stem Cell Institute and the Broad Institute, a Swiss National Science Foundation fellowship (MS), a Human Frontier Science Program fellowship (DC), an ERC Advanced grant (ERC-AdG 294325-GeneNoiseControl) and a Nederlandse Organisatie voor Wetenschappelijk Onderzoek (NWO) Vici award (AvO).

## Supplementary Data

All sequences and derived gene expression data have been deposited in the NCBI GEO database under accession number GSE53638.

## References

1. Shapiro, E., Biezuner, T. & Linnarsson, S. Single-cell sequencing-based technologies will revolutionize whole-organism science. Nat. Rev. Genet. 1–13 (2013). doi:10.1038/nrg3542

2. Islam, S. et al. Characterization of the single-cell transcriptional landscape by highly multiplex RNA-seq. Genome Res. 21, 1160–7 (2011).

3. Islam, S. et al. Highly multiplexed and strand-specific single-cell RNA 5′ end sequencing. Nat. Protoc. 7, 813–28 (2012).

4. Hashimshony, T., Wagner, F., Sher, N. & Yanai, I. CEL-Seq: single-cell RNA-Seq by multiplexed linear amplification. Cell Rep. 2, 666–73 (2012).

5. Kapteyn, J., He, R., McDowell, E. T. & Gang, D. R. Incorporation of non-natural nucleotides into template-switching oligonucleotides reduces background and improves cDNA synthesis from very small RNA samples. BMC Genomics 11, 413 (2010).

6. Gertz, J. et al. Transposase mediated construction of RNA-seq libraries. Genome Res. 22, 134–41 (2012).

7. Kivioja, T. et al. Counting absolute numbers of molecules using unique molecular identifiers. Nat. Methods 9, 72–4 (2012).

8. Jaitin, D. A. et al. Massively parallel single-cell RNA-seq for marker-free decomposition of tissues into cell types. Science 343, 776–9 (2014).

9. Picelli, S. et al. Smart-seq2 for sensitive full-length transcriptome profiling in single cells. Nat. Methods 10, 1096–8 (2013).

10. Aust, L. et al. Yield of human adipose-derived adult stem cells from liposuction aspirates. Cytotherapy 6, 7–14 (2004).

11. Rosen, E. D. & MacDougald, O. a. Adipocyte differentiation from the inside out. Nat. Rev. Mol. Cell Biol. 7, 885–96 (2006).

12. Cristancho, A. G. & Lazar, M. a. Forming functional fat: a growing understanding of adipocyte differentiation. Nat. Rev. Mol. Cell Biol. 12, 722–34 (2011).

13. Green, H. & Meuth, M. An established pre-adipose cell line and its differentiation in culture. Cell 3, 127–33 (1974).

14. Mikkelsen, T. S. et al. Comparative epigenomic analysis of murine and human adipogenesis. Cell 143, 156–69 (2010).

15. Raj, A., van den Bogaard, P., Rifkin, S. A., van Oudenaarden, A. & Tyagi, S. Imaging individual mRNA molecules using multiple singly labeled probes. Nat. Methods 5, 877–9 (2008).

16. Brunet, J.-P., Tamayo, P., Golub, T. R. & Mesirov, J. P. Metagenes and molecular pattern discovery using matrix factorization. Proc. Natl. Acad. Sci. U. S. A. 101, 4164–9 (2004).

17. Subramanian, A. et al. Gene set enrichment analysis: a knowledge-based approach for interpreting genome-wide expression profiles. Proc. Natl. Acad. Sci. U. S. A. 102, 15545–50 (2005).

18. Eguchi, J. et al. Interferon regulatory factors are transcriptional regulators of adipogenesis. Cell Metab. 7, 86–94 (2008).

19. Eguchi, J. et al. Transcriptional control of adipose lipid handling by IRF4. Cell Metab. 13, 249–59 (2011).

20. Pietiläinen, K. H. et al. Global transcript profiles of fat in monozygotic twins discordant for BMI: pathways behind acquired obesity. PLoS Med. 5, e51 (2008).

21. Lee, Y. H. et al. Microarray profiling of isolated abdominal subcutaneous adipocytes from obese vs non-obese Pima Indians: increased expression of inflammation-related genes. Diabetologia 48, 1776–83 (2005).

22. Mutch, D. & Tordjman, J. Needle and surgical biopsy techniques differentially affect adipose tissue gene expression profiles. Am. J. Clin. Nutr. 89, 51–57 (2009).

23. Toubal, A., Treuter, E., Clément, K. & Venteclef, N. Genomic and epigenomic regulation of adipose tissue inflammation in obesity. Trends Endocrinol. Metab. 24, 625–34 (2013).

